# Cellular hnRNPA0 limits HIV-1 production by interference with LTR-activity and programmed ribosomal frameshifting

**DOI:** 10.1101/2023.08.08.552510

**Authors:** Fabian Roesmann, Helene Sertznig, Katleen Klaassen, Alexander Wilhelm, Delia Heininger, Carina Elsner, Mario Santiago, Stefan Esser, Kathrin Sutter, Ulf Dittmer, Marek Widera

## Abstract

The interplay between host factors and viral components has a profound impact on the viral replication efficiency and fitness. Heterogeneous nuclear ribonucleoproteins (hnRNPs), in particular members of the subfamily A/B, have been broadly studied as HIV-1 host dependency factors, however, the least related member hnRNPA0 has so far not been functionally studied in its potential role affecting viral replication.

In this study, we revealed that hnRNPA0 overexpression in HEK293T cells significantly reduced HIV-1 long terminal repeat (LTR) activity up to 3.5-fold, leading to a significant decrease in total viral mRNA (5.5-fold) and protein levels (3-fold). Conversely, knockdown of hnRNPA0 enhanced LTR activity, suggesting its negative regulatory role in viral gene expression. Moreover, the splicing pattern of HIV-1 remained largely unaffected by altered hnRNPA0 levels indicating changes in viral mRNA expression predominantly occurred at the transcriptional level. Moreover, hnRNPA0 overexpression was found to significantly reduce the programmed ribosomal frameshift efficiency of HIV-1, resulting in a shift in the HIV-1 p55/p15 ratio, compromising viral fitness. Synergistic inhibition of LTR activity and thus reduced viral mRNA transcription and impaired ribosomal frameshifting efficiency, which is important for viral infectivity, were detrimental to HIV-1 replication. Additionally, our study revealed that hnRNPA0 levels were lower in therapy naïve HIV-1-infected individuals compared to healthy controls and temporarily repressed after IFN-I treatment in HIV-1 target cells.

Our findings highlight the significant role of hnRNPA0 in HIV-1 replication and suggests that its IFN-I regulated expression levels are decisive for viral fitness.

**Importance:** RNA binding proteins, in particular heterogeneous nuclear ribonucleoproteins (hnRNPs) have been extensively studied as host dependency factors for HIV-1 since they are involved in multiple cellular gene expression processes. However, the functional role of hnRNPA0, the least related member of the hnRNPA/B family, and its potential impact on viral replication remains unclear. For the first time, our findings demonstrate the significance of hnRNPA0 in restricting viral replication efficiency. We demonstrate that hnRNPA0 plays a pleiotropic role in limiting viral replication being a negative regulator of viral transcription and significantly impairing ribosomal frameshifting. Our study also revealed hnRNPA0 as an IFN-regulated host factor that is temporarily repressed after IFN-I treatment in HIV-1 target cells and lower expressed in therapy-naïve HIV-1-infected individuals compared to healthy controls. Understanding the mode of action between hnRNPA0 and HIV-1 might help to identify novel therapeutically strategies against HIV-1 and other viruses.

## Introduction

The human immunodeficiency virus type 1 (HIV-1) is the causative agent of the acquired immunodeficiency syndrome (AIDS). HIV-1 is highly adapted to the human host and can exploit various host factors (1–3) and hijack essential cellular processes for its own replication. After infection of a susceptible host cell, predominantly CD4^+^ T-cells and macrophages, the single-stranded 9.7kb long RNA is reverse transcribed into double-stranded DNA, which then is irreversibly integrated into the genome of the host cell (4, 5). HIV-1 uses a variety of cellular mechanisms to express its complex genome. The HIV-1 replication strategy includes alternative splicing of its own pre-mRNA (6) by a diverse network of splicing-regulatory-elements (SREs) acting in *cis-* and cellular RNA-binding proteins (RBPs) in *trans*-binding SREs (7). By binding the *cis-*regulatory elements, heterogeneous nuclear ribonucleoproteins (hnRNPs) and serine/arginine-rich-splicing factors (SRSFs) play a crucial role in alternative splicing and are thus essential for HIV-1. To further expand the genetic repertoire, HIV-1 induces a programmed ribosomal frameshift (PRF) to translate two of its three reading frames, resulting either in synthesis of the viral structural proteins Gag (i.a. matrix, capsid, nucleocapsid), or upon -1PRF, generation of the viral enzymes Gag-Pol (i.a. protease, reverse transcriptase, integrase) (8).

HnRNPs are ubiquitously expressed in various cell types and tissues. They are able to bind distinct conserved RNA motifs and by doing so might influence the stability, localization, processing and the functionality of cellular and viral RNAs (9). hnRNPA1 and A2/B1 have been extensively studied in many aspects, also several publications focusing on HIV-1 are published for these two proteins (10–16) and hnRNPA3 (7, 17–20).

With 305 aa hnRNPA0 is the shortest member of the hnRNPA/B family and has comparably received little attention. HnRNPA0 preferably binds to adenylate-uridylate (AU)-rich elements (AREs), which are commonly located at 3’-untranslated regions (UTRs) of mRNAs (21–24). The consensus motif of the AREs bound by hnRNPA0 is the pentamer AUUUA (23), which is notably also bound by hnRNPA1 (25). Mechanistically, it has been shown that hnRNPA0 together with hnRNPA2B1 and ELAV Like RNA Binding Protein 1 (ELAVL1) binds to the 5’UTR of cellular *AXIIR* mRNA and by cooperating with upstream open-reading-frames inhibits translation of AXIIR, a receptor important for annexin II-mediated osteoclastogenesis (26, 27). In multiple cases, changes in hnRNPA0 sequence composition, expression and modifications were associated with multiple common cancers like prostate, breast, or colon, but also with several uncommon cancers (28). In particular, hnRNPA0 might also function as a biomarker (29) for cancers like hereditary colorectal cancer (30).

It was shown that tumor-specific phosphorylation of hnRNPA0 inhibited apoptosis though the promotion of the G2/M phase maintenance in colorectal cancer cells (31) indicating a role of hnRNPA0 in apoptosis and cell cycle regulation. By binding to the AREs of the respective mRNAs, hnRNPA0 is also involved in the processing of immunomodulatory mRNAs encoding tumor necrosis factor alpha (TNF-alpha), cyclooxygenase 2 (Cox-2) and macrophage inflammatory protein-2 (MIP-2) (23). Furthermore, mutation analysis revealed that hnRNPA0 is involved in the ERK/MAPK signaling pathways, which also play a crucial role in regulating processes such as cell growth, cell differentiation, cell survival, cell migration, and cell division (28).

Whether hnRNPA0 might also affect viral replication, is so far unclear and needs further investigation. Since other members of the hnRNPA/B subfamily are described to modulate gene expression of cellular and viral genes, in this manuscript we investigated the impact of hnRNPA0 on HIV-1 and focused on post integration steps.

As overexpression resulted in severe impairments of HIV-1 LTR transcription, -1PRF, and virus production, our findings add another role to the diverse functions of hnRNPA0. In addition, this work highlights the so far undescribed role of hnRNPA0 as a strong antiviral IFN-regulated cellular effector molecule in viral infections.

## Results

### hnRNPA0 overexpression reduces HIV-1 particle production and infectivity

The hnRNPA/B family consists of four members that are hnRNPA0, A1, A2B1, and A3. Point-accepted mutation analysis (PAM; Fig. 1a) revealed that hnRNPA0 is quite distinct from the other hnRNPA/B members, despite carrying all main characteristics, two RNA-recognition-motifs (RRM) and an unstructured glycine-rich region (Fig. 1b) (32). In particular, the C-terminally located glycine-rich region of hnRNPA0 differs compared to the other family members (Supp.Fig.3). HnRNPA0 exhibited the highest similarity to hnRNPA2B1 (score: 0.577), while hnRNPA1, A2B1, and A3 had an average PAM score of 0.264, revealing a higher degree of similarity. While several interactions between HIV-1 and the hnRNPA/B family have been reported so far, little is known about the role of hnRNPA0 in the context of retroviruses.

**Figure 1.**
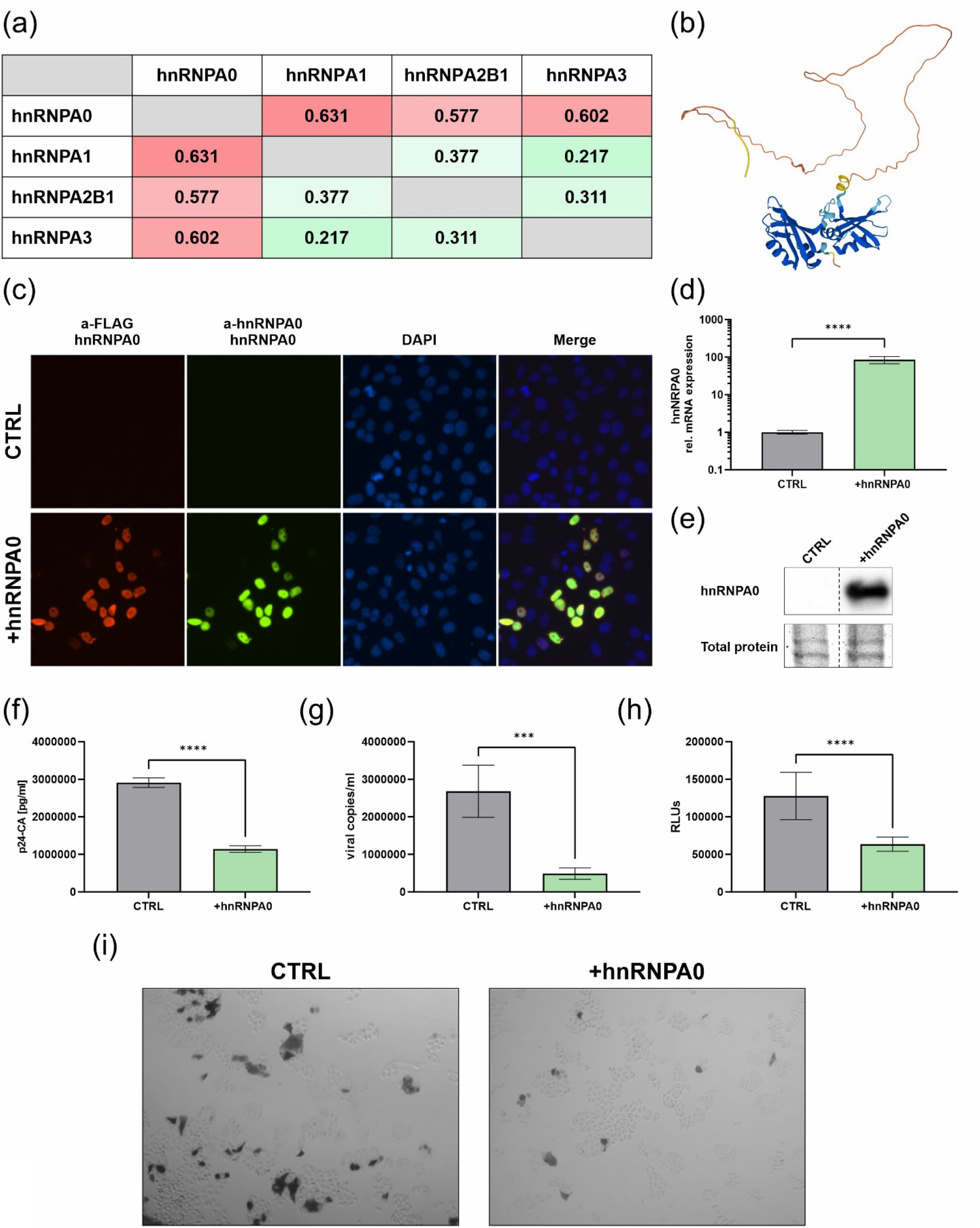
Overexpression of hnRNPA0 limits production of infectious HIV-1 particles. (a) Point accepted mutation 120 matrix (PAM 120) of the hnRNPA/B family. For each hnRNP the canonical sequence was chosen from the Uniprot database (33). Entry IDs: hnRNPA0: Q13151, hnRNPA1: P09651, hnRNPA2B1: P22626, hnRNPA3: P51991. **(b)** Protein structure of hnRNPA0 was predicted using Alphafold (34, 35). **(c-e)** Vero **(c,e)** or HEK293T **(d)** cells were transfected with an hnRNPA0 expressing vector. **(c)** 24 h post transfection cells were fixed using 3% formaldehyde for 10 min at RT and permeabilized using 0.1% Triton X-100 for 10 min. Cells were rinsed twice with PBS and unspecific binding sites were blocked using 2% BSA for 20 min before samples were incubated for 1 h with antibodies against cellular and FLAG-tagged hnRNPA0 following a washing step and an 1 h incubation with secondary antibodies. Nuclei were stained using DAPI **(d)** 24 h post transfection RNA was isolated and expression levels were analyzed via RT-qPCR. **(e)** 24 h post transfection proteins were isolated and Western blotting was performed to analyze the protein amount using an antibody against hnRNPA0. One representative Western blot of three independent experiments is shown. **(f-i)** HEK293T were transfected with the proviral molecular clone NL4-3 and an hnRNPA0 expressing vector. 48 h post transfection viral supernatant was harvested and the **(f)** particle production was analyzed via p24-Capsid-ELISA, the **(g)** viral copy numbers were analyzed using RT-qPCR and the **(h,i)** presence of infectious particles in the cell culture supernatant was analyzed using TZM-bl reporter cells **(h)** luciferase assay and **(i)** X-Gal staining. Mean (+SD) of four independent experiments is shown for (d,f,g,h) Unpaired two-tailed t-tests were performed to determine statistical significance (*p<0.05, **p<0.01, ***p<0.001, ****p<0.0001).

*In silico* mapping of hnRNPA0 binding motifs to the HIV-1 reference sequence NL4-3 revealed the presence of multiple hnRNPA0 binding sites, in particular in the Gag-Pol, and splice donor 4 (D4) region (Supp.Fig.1). To analyze whether the putative binding sites of hnRNPA0 in the HIV-1 genome might indicate a possible regulatory role for hnRNPA0 in viral replication, we generated an hnRNPA0 expression vector, evaluated expression levels and localization of the flag-tagged protein (Fig. 1c-e), and proceeded to co-transfect HEK293T cells with the proviral molecular HIV-1 clone NL4-3. 48 h post transfection the viral supernatant was harvested and used for subsequent analysis (Fig. 1f-i). Upon elevated hnRNPA0 levels (85.4-fold; p<0.0001) we observed a significant decrease in p24-CA production (3-fold; p<0.0001), viral copy numbers (> 5-fold; p=0.0008) in the supernatant and production of infectious particles (2-fold; p<0.0001). These data indicate, that high levels of hnRNPA0 limit HIV-1 replication and possess a potent antiviral activity.

### hnRNPA0 modulates HIV-1 Tat mediated LTR activity

Next, we investigated the potential influence of hnRNPA0 on viral replication steps as hnRNPs are responsible for the regulation of alternative splicing in HIV-1. We assumed, that hnRNPA0 may decrease viral replication by causing aberrant splice site usage, resulting in insufficient or unbalanced mRNA transcript amounts (7). Hence, RT-qPCR was conducted to assess viral splicing under varying hnRNPA0 levels – overexpression and knockdown conditions (Fig. 2). Viral splicing pattern was analyzed using semi-quantitative RT-PCR (Fig. 2a,g) with primers targeting specified sites as outlined in (36). Upon high hnRNPA0 levels we observed an increase in *vif1*, *vpr1, tat2* (tat-mRNA-class) and *nef1* (2kb-class). *Vpr3, tat5* (4kb-class) and *tat1* (2kb- class) levels were slightly decreased. RT-qPCR revealed a strong reduction in total viral mRNA, measured by primer pairs targeting the initial and terminal HIV-1 exons 1 (2.93-fold; p=0.0089) and 7 (4.56-fold; p<0.0001), indicating an inhibitory effect on HIV-1 LTR activity. By using intron 1 spanning primers and those covering the 2kb class specific exon junction D4-A7, we observed a trend towards more unspliced and significantly less multiply spliced mRNAs (0.8-fold; p=0.03) (Fig. 2c). Exon 2 containing transcripts, however, were detected more frequently (p=0.0088), while those harboring exon 3 were significantly less detected (p=0.0388). Although *in silico* mapping of hnRNPA0 to the NL4-3 genome revealed a strong binding affinity towards D4 (3.97 Z-score (Supp.Fig.1), we did not observe a major effect on alternative spice site usage under high hnRNPA0 conditions. When using primers spanning the exonic HIV-1 D1-A1 splice site junction including the *vif* intron, we observed a 1.8-fold (p=0.0424) increase in the mRNA coding for the essential accessory protein Vif, which counteracts the host restriction factor APOBEC3G (37) and the cytosolic DNA sensor STING (38). Further usage of primers specific for the HIV-1 D1-A2 splice site junction and the *vpr* intron, revealed a slight increase in the splicing efficiency of *Vpr*, a pleiotropic accessory protein necessary for replication in certain non-permissive cell lines like dendritic cells (39).

**Figure 2.**
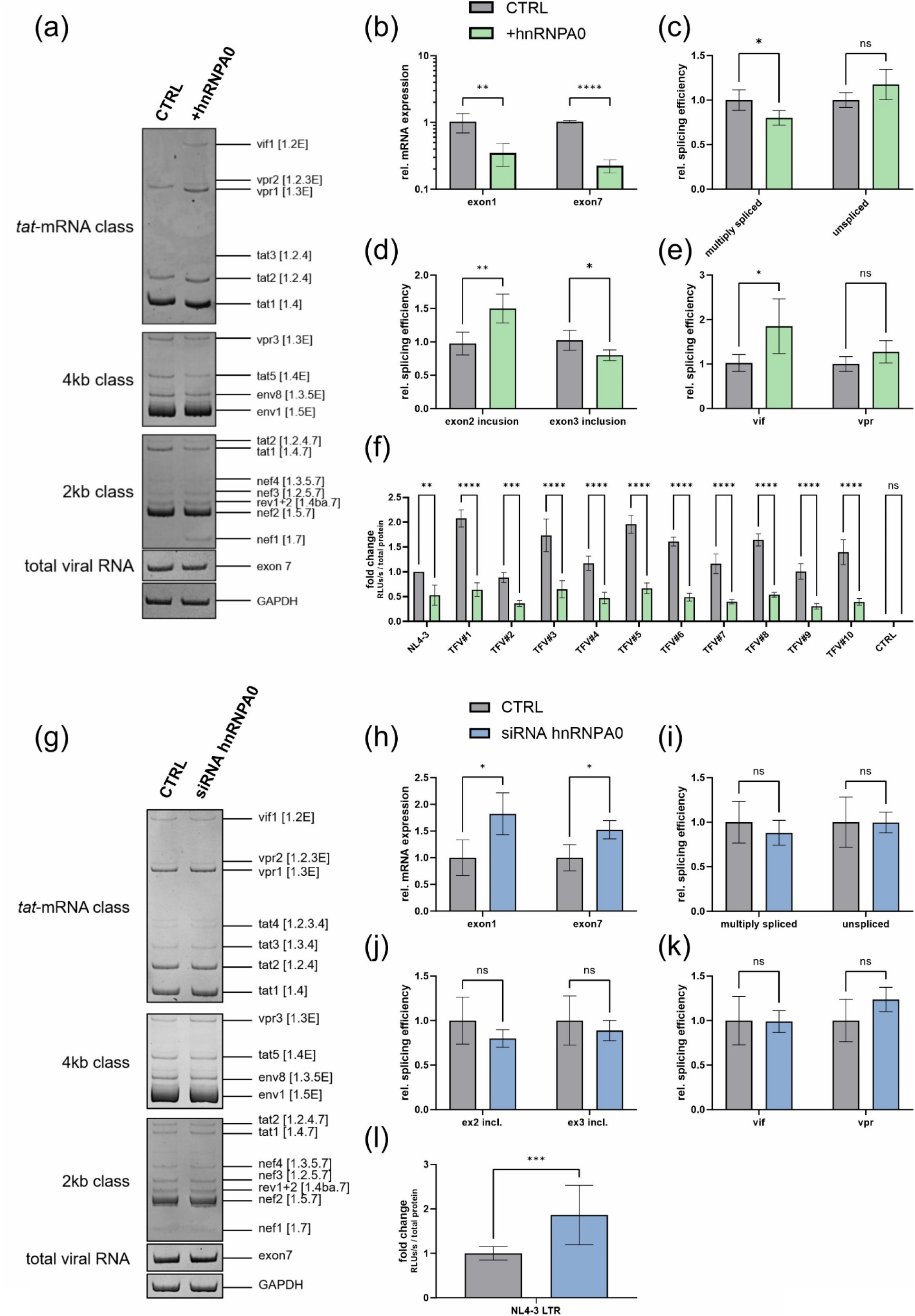
HIV-1 splicing pattern and LTR-activity at high and low hnRNPA0 conditions. **(a-e)** HEK293T cells were transfected with a plasmid coding for the proviral clone NL4-3 (pNL4-3) and an expression vector encoding hnRNPA0 or an empty vector control (pcDNA3.1(+)). 48 h post transfection RNA was isolated and subjected to further analysis. **(g-k)** HEK293T cells were transfected with pNL4-3 as well as siRNA against hnRNPA0 or an off-target control. 72 h post transfection cells were lysed, RNA isolated and RT-qPCR was performed to evaluate expression levels. **(a)** RT-PCR was performed using primer pairs covering viral mRNA isoforms of the 2kb, 4kb and tat-mRNA- class (described in (36). Primers covering HIV-1 exon 7 containing transcripts were used for normalization of whole-viral-mRNA and cellular GAPDH was included as loading control. The HIV-1 transcript isoforms are labelled according to (40). The amplified PCR products were separated on a 12% non-denaturing polyacrylamide gel. **(b)** expression levels of exon 1 and exon 7 containing mRNAs (total viral mRNA) were normalized to GAPDH. **(c-e)** Expression levels of **(c)** multiply spliced and unspliced mRNAs, **(d)** exon 2 and exon 3 containing mRNAs, **(e)** *vif* and *vpr* were normalized to exon 1 and exon 7 containing mRNAs (total viral mRNA). **(f)** Vero cells were transfected with plasmids coding for hnRNPA0 (pcDNA-FLAG-NLS-hnRNPA0), Tat (SVcTat) and a Firefly luciferase reporter plasmid with LTR sequences from the proviral clone NL4-3 sequences obtained from transmitted founder viruses (TFV) from patient samples (TFV#1-10) (41, 42) provided by Dr. John C. Kappes. A vector (pTA_Luc) expressing only the Firefly luciferase was used as control. 24h post transfection the cells were lysed and luciferase-based reporter assays were performed. The relative light units (RLUs) were normalized to the protein amount analyzed via Bradford assay. **(l)** A549 LTR Luc-PEST reporter cells were transfected with a plasmid encoding the Tat protein (SVcTat) as well as siRNA against hnRNPA0 or an off-target control. 48 h post transfection cells were lysed and luciferase reporter assays were performed. The RLUs were normalized to the protein amount analyzed via Bradford assay. The reporter cell line was previously generated using the Sleeping Beauty system (43). The PEST sequence fused to the Firefly luciferase causes a rapid degradation of the protein (44). Mean (+SD) of four biological replicates for (b-e & h-k), three replicates for (f) and twelve independent replicates from two biologically independent experiments for (l). Unpaired two-tailed t-tests were calculated to determine whether the difference between sample groups reached the level of statistical significance (*p<0.05, **p<0.01, ***p<0.001, ****p<0.0001 and ns, not significant) for **(f)** two-way ANOVA with Dunnett post-hoc test was performed.

To investigate whether the decrease of total viral mRNA, was due to an influence of hnRNPA0 on the LTR activity, we performed luciferase-based reporter assays. Cells were co-transfected with Tat-transactivated LTR-luciferase reporter constructs, and the hnRNPA0 expression vector. 24 h post transfection cells were lysed and the luciferase activity was measured. In addition to the LTR sequence of the laboratory strain NL4-3 we tested LTR sequences derived from ten HIV-1 transmitted-founder viruses (TFV, Fig. 2f). We observed an almost 2-fold reduction of the NL4-3 LTR (p=0.0028) activity, while TFV LTR activities were repressed to a greater degree.

To investigate whether hnRNPA0 knockdown would lead to an opposite effect, we co- transfected cells with a plasmid encoding the proviral clone NL4-3 and siRNA against hnRNPA0. Despite a small increase in *vpr3* (4kb-class; Fig. 2g) we did not observe any changes in the viral splicing pattern under depleted hnRNPA0 levels using the semi-quantitative approach. Furthermore, we quantified the total viral mRNA (reflected by exon 1 and exon 7 expression). In contrast to high hnRNPA0 levels, we observed an increase in HIV-1 exon 1 (1.8-fold; p=0.0187) and exon 7 (1.5-fold; p=0.0124) containing mRNAs upon hnRNPA0 knockdown. RT-qPCR revealed no significant changes in the viral splicing pattern (Fig. 2g-h). Noteworthy, we did not observe a decrease in *vif* splicing efficiency.

To emphasize this finding, we monitored the activity of the NL4-3 LTR via luciferase reporter cells and detected a 1.86-fold increase in LTR activity (p=0.0003) under depleted hnRNPA0 levels (Fig. 2l).

### hnRNPA0 affects HIV-1 protein levels by modulating LTR activity and frameshifting

Next, we addressed whether the observed transcriptional and post-transcriptional effects would result in altered protein levels (Fig. 3). At first, we confirmed the overexpression of hnRNPA0 in the transfected cells and observed an almost 10-fold increase (p=0.0003) in hnRNPA0 protein amounts (Fig. 3a-b). The Vif and Vpr protein levels reflected the earlier splicing-efficiency observations with Vif (1.89-fold; p=0.0319) and Vpr (2.22-fold; p=0.023) levels being decreased. Furthermore, we observed a strong repression in p24 levels (3.03-fold; p=0.0009). Thus, we observed an overall significant reduction of essential HIV-1 proteins upon elevated hnRNPA0 levels.

**Figure 3.**
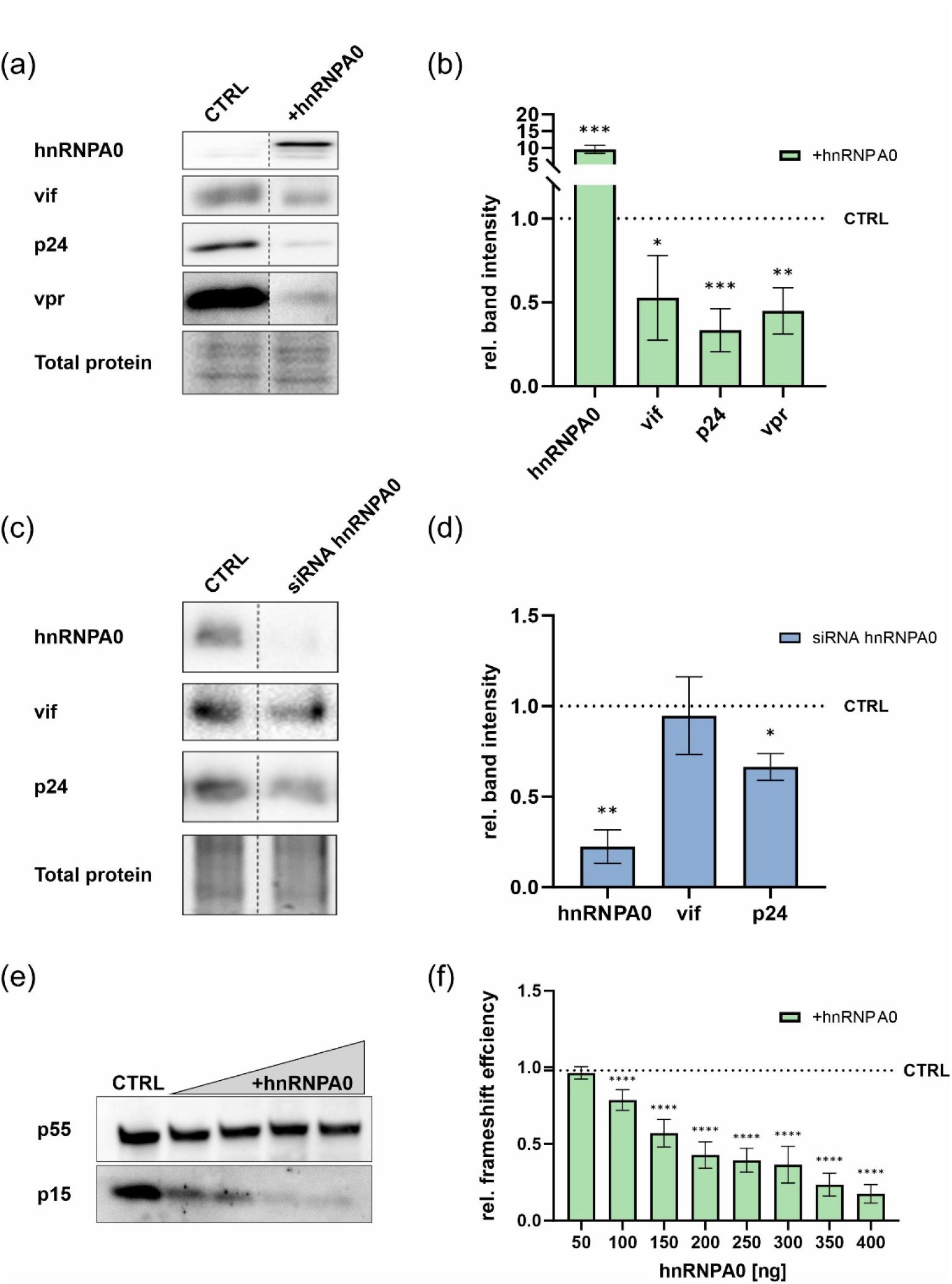
HIV-1 protein levels are modulated upon hnRNPA0 overexpression or knockdown. HEK293T cells were transfected with a plasmid encoding the proviral clone NL4-3 and an expression vector coding for hnRNPA0. 48 h post transfection cells were lysed and the protein amounts were analyzed via Western blotting using the antibodies listed in (Tab.1). Trichloroethanol was used to stain total protein amounts, which were further used for normalization. **(a)** Representative Western blot of four independent replicates for the overexpression of hnRNPA0 quantified in **(b)**. For improved comparability, the samples were repositioned adjacently after imaging, as denoted by the dotted line. The depicted samples underwent processing on the same nitrocellulose membrane. **(c-d)** HEK293T cells were transfected with pNL4-3 as well as siRNA against hnRNPA0 or an off-target control. 72h post transfection cells were lysed and Western blotting was performed to evaluate the protein amounts. **(c)** Representative Western blot of four independent replicates of the quantification shown in **(d)**. **(e)** HEK293T cells were transfected with pNL4-3 and increasing amounts of a plasmid encoding hnRNPA0 (250, 500, 1000, 1500 ng). 48 h post transfection cells were lysed, proteins were separated via PAGE and analyzed via immunoblotting using an antibody targeting p15. One representative Western blot of three independent experiments is shown **(f)** HEK293T cells were transfected with the indicated amounts of a plasmid encoding hnRNPA0. 6h post transfection a second transfection was performed using luciferase reporter plasmids including the HIV-1 frameshift site. 24h post second transfection the cells were harvested and the Firefly to Renilla luciferase activity ratio was measured via luciferase reporter assay. In the luciferase reporter the Renilla luciferase is positioned in-frame, facilitating translation during ribosomal scanning of the RNA, while the Firefly luciferase is placed in the -1-frame, yielding a functional polyprotein only upon occurrence of the -1 frameshift. Mean (+SD) of three biologically independent experiments with three replicates each is shown. For 350 and 400 ng one experiment with three independent replicates was performed. Unpaired two-tailed t-tests were calculated to determine whether the difference between the sample groups reached the level of statistical significance (*p<0.05, **p<0.01, ***p<0.001, ****p<0.0001 and ns, not significant), for **(f)** Mixed-effects analyses followed by Dunnett post-hoc test were performed.

Upon knockdown, the protein levels of hnRNPA0 were decreased 4.46-fold (p=0.0011) (Fig. 3d). Although we previously observed enhanced transcriptional activity under depleted hnRNPA0 levels, the protein levels of p24 were significantly reduced (1.51-fold; p=0,0009), and Vif levels were also not elevated compared to the control. This led to the assumption, that hnRNPA0 might not only regulate LTR-activity but might also be involved in late post-transcriptional steps.

Indeed, we noticed a shift between HIV-1 p15 and p55 upon high hnRNPA0 levels. A shift in the p15/p55-ratio would indicate an effect on the viral-frameshift efficiency, as both genes are translated from the same unspliced mRNA. Indeed, we observed a dose-dependent inhibition of PRF by a reduced p15/p55 ratio dropping from 1.18 to 0.53 (Fig. 3e). To proof this hypothesis, we used a luciferase reporter plasmid encoding a Renilla luciferase, the viral frameshift site of HIV-1 and a Firefly luciferase (Fig. 3f). Supporting the previous results, we observed a dose-dependent reduction of -1PRF efficiency upon increased hnRNPA0 levels. Thus, we concluded that hnRNPA0 is not only capable of modulating the HIV-1 LTR-activity, but it is also able to modify frameshifting efficiency modulating the viral replication.

With the contradictory results on mRNA and protein levels under hnRNPA0 knockdown conditions we performed end-point readouts of the previously performed knockdown experiments.

### Knockdown of hnRNPA0 enhance HIV-1 particle production and infectivity

We observed significantly more particles in the viral supernatant under depleted hnRNPA0 conditions (Fig. 4a). Concomitantly, viral copies in the supernatant were also significantly increased (1.12-fold; p=0.0117; Fig. 4b). To proof whether low hnRNPA0-levels might facilitate viral replication even in the presence of host restriction factors, we transfected A3G expressing cells with an anti-hnRNPA0 siRNA and a NL4 3 encoding plasmid (Fig. 4a). The subsequent luciferase assay revealed increased infectivity of the supernatants from cells low and intermediate expressing APOBEC3G (Fig. 4c). At higher APOBEC3G levels we did not observe significant increased infectivity. Thus, depleted hnRNPA0 levels facilitate HIV-1 infectivity even in APOBEC3G expressing cells.

**Figure 4.**
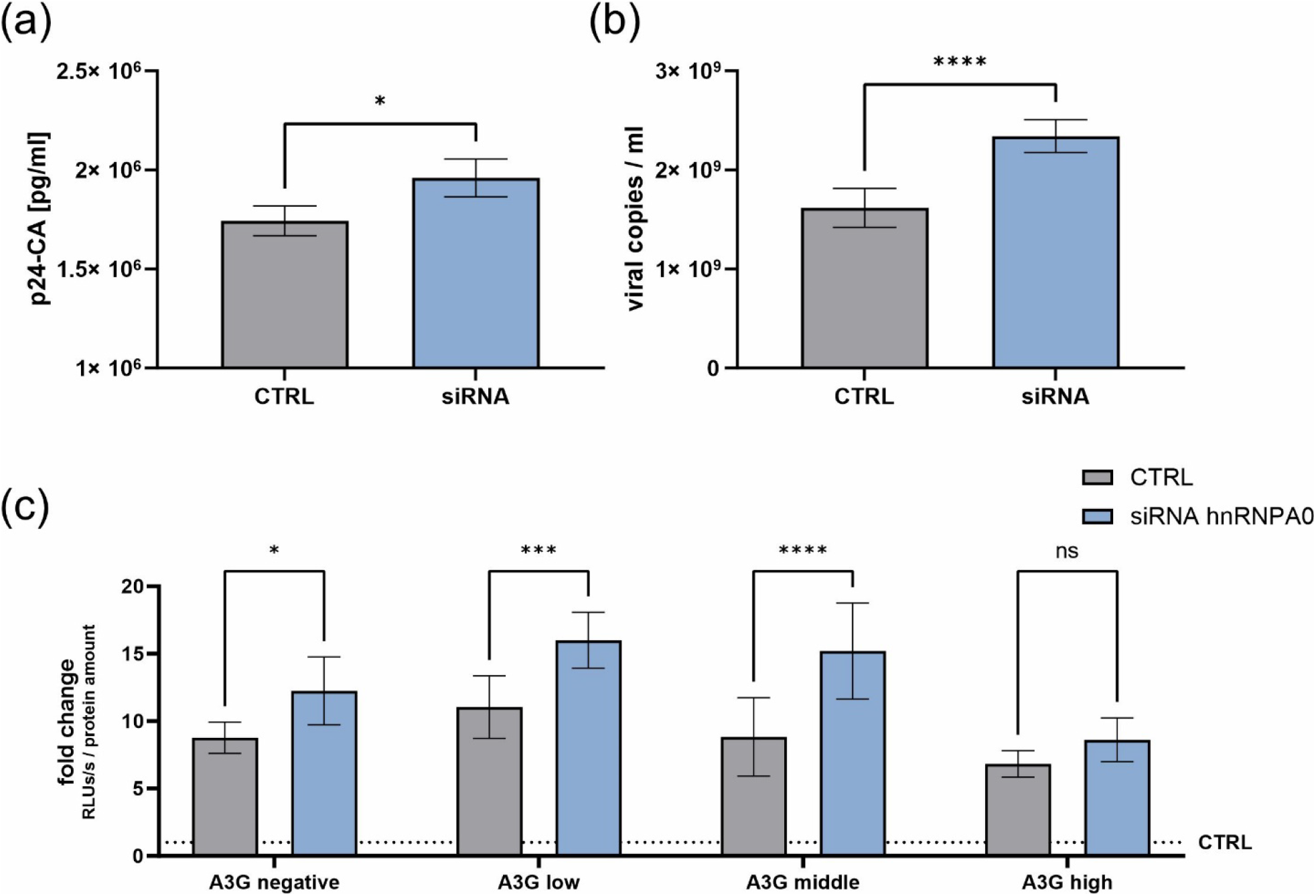
Knockdown of hnRNPA0 elevates viral particle production, copy numbers and infectivity. HEK293T cells were transfected with a plasmid coding for the proviral clone NL4-3 and siRNA against hnRNPA0 and an off-target control. 72 h post transfection the supernatant was harvested and used for subsequent experiments. Particle production was analyzed via **(a)** p24-Capsid-ELISA. **(b)** RNA of the viral supernatant was isolated and RT-qPCR was performed to evaluate viral copy numbers. **(c)** Viral supernatant was used to infect TZM-bl reporter cells. 48 h post infection TZM-bl cells were lysed and the luciferase activity was measured. The relative light units were normalized to the total protein amount analyzed via Bradford assay. Mean (+SD) is shown of four independent replicates of (a), six of (b) and four of (c) with two technical replicates each. Unpaired two-tailed t-tests were calculated to determine whether the difference between the sample groups reached the level of statistical significance (*p<0.05, **p<0.01, ***p<0.001, ****p<0.0001 and ns, not significant) for (c) the groups were compared by two-way ANOVA with Bonferroni post hoc test.

### hnRNPA0 expression levels are regulated by interferons

Interferons play a critical role in limiting HIV-1 replication by inducing an antiviral state in the infected and bystander cells (45, 46). In a previous study, we could demonstrate a significant deregulation of the RNA binding protein SRSF1 in IFN treated macrophage-like THP-1 cells. Since we found that high levels of hnRNPA0 were detrimental for viral replication but low levels of hnRNPA0 seem to boost the infectivity of HIV-1, we were interested whether hnRNPA0 levels might be regulated upon IFN stimulations. Hence, comparable settings were used and phorbol 12-myristate 13-acetate (PMA)-differentiated THP-1 cells were subjected to infection experiments using the CCR5-tropic HIV-1 NL4-3 AD8 strain (47). Additionally, we treated the cells 16 h post infection with IFNα14, as this was reported to be the most potent IFN-I against HIV-1 (48, 49) and was previously shown to strongly repress SRSF1 expression (36). To investigate whether hnRNPA0 might be specifically regulated by exogenous IFNs we added a Jak1-2 inhibitor Ruxolitinib, preventing IFN signaling (Fig. 5). At the indicated time points total cellular RNA was isolated and expression levels were analyzed via RT-qPCR.

**Figure 5.**
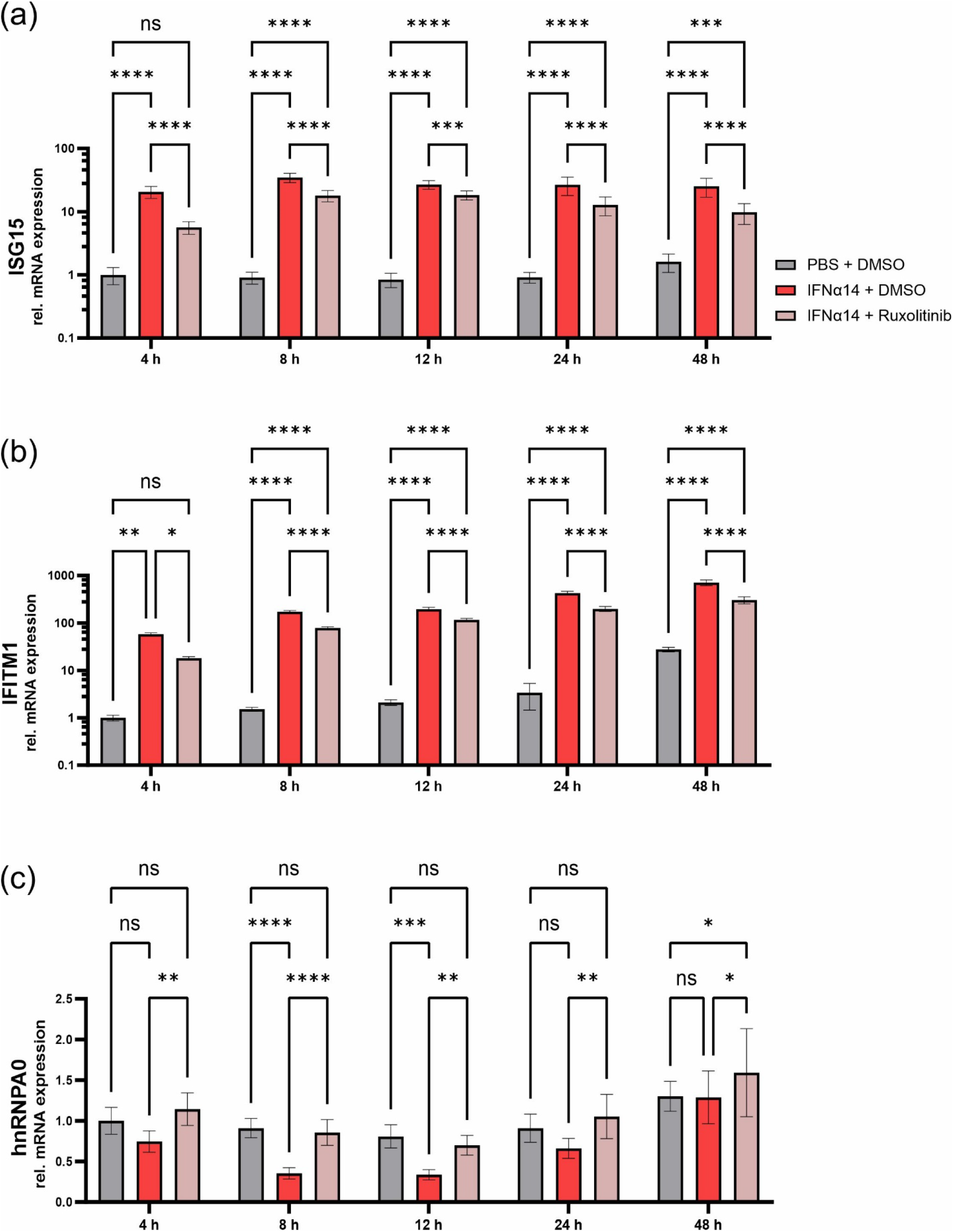
Interferon signaling regulates hnRNPA0 mRNA expression. Differentiated THP-1 macrophages were treated with 1 µM Ruxolitinib or a DMSO solvent control 1 h before inoculation of the viral clone NL4-3 AD8 (MOI 1). 16 h post infection cells were washed and medium containing 10 ng/ml IFNα14 or PBS and 1 µM Ruxolitinib or DMSO was added. At the indicated time points post treatment, cells were lysed and further subjected to RNA isolation. RT-qPCR was performed to monitor mRNA expression levels of **(a)** ISG15 **(b)** IFITM1 and **(c)** hnRNPA0. Mean (+SD) of two biologically independent experiments were performed with 4 biological replicates each. The groups were compared with two-way repeated-measures ANOVA with Tukey’s post-hoc test to determine whether the difference between the sample groups reached the level of statistical significance (*p<0.05, **p<0.01, ***p<0.001, ****p<0.0001 and ns, not significant).

To monitor IFN-stimulation we additionally measured *ISG15* and *IFITM1* (Fig. 5a-b). A significant increase in *ISG15* and *IFITM1* mRNA expression, was observed upon IFN-treatment throughout all time points. Of note, even though the Ruxolitinib-treated cells showed elevated *ISG15* and *IFITM1* mRNA expression, the expression levels remained below the samples lacking the JAK1-2 inhibitor. In agreement with previous studies, the induction of ISG15 and IFIMT1 despite inhibition of JAK1/2 could be explained by combinatory effects of the IFN treatment and viral sensing via TLRs (50–52). While the *hnRNPA0* mRNA expression levels of the Ruxolitinib-treated cells were unaffected, we observed a reduction of *hnRNPA0* mRNA expression in the solely IFN-treated cells, already at 4h post treatment. After 8h (2.56-fold; p<0.0001) the mRNA expression was significantly decreased and further waned until 12h post treatment but was fully restored after 48h. These results indicated that hnRNPA0 is regulated by JAK1-2 dependent IFNs signaling.

To further evaluate IFN-mediated regulation of hnRNPA0, we treated both HIV-1 target cell types using cell lines THP-1 (Fig. 6a-d) and Jurkat (Fig. 6e-h) with IFN-I. We additionally included IFNα2 since it is used in clinical treatment of chronic viral infections like hepatitis-B-virus (53). *ISG15* was again used as surrogate marker for IFN induced ISG induction. Similar to the previous experiment, the mRNA expression of *hnRNPA0* was significantly reduced after 8h, while the strongest repression was observed at 12h post treatment. After 24h the mRNA expression increased again until it was almost recovered after 48h. A similar, although stronger and longer lasting effect, was observed when treating THP-1 cells with IFNα14. In Jurkat T-cells *ISG15* expression after IFNα2 treatment was strongly induced 4h and 8h post stimulation, but the levels rapidly declined after 12h (Fig. 6e). The *ISG15* expression in IFNα14-treated Jurkat cells was high and long lasting, and comparable to IFN treated THP-1 cells. While no alteration in *hnRNPA0* expression was observed in IFNα2-treated Jurkat cells, an initial tendency for lower *hnRNPA0* mRNA levels 8-24h post treatment with IFNα14 was observed. Strikingly, a significant (p=0.0233) increase in mRNA levels was measured 48h post stimulation. As we observed the strongest reduction in *hnRNPA0* mRNA levels in IFNα14-treated THP-1 cells, we also confirmed the decrease in protein amount under these conditions using Western blot analysis (Fig. 6I-j).

**Figure 6.**
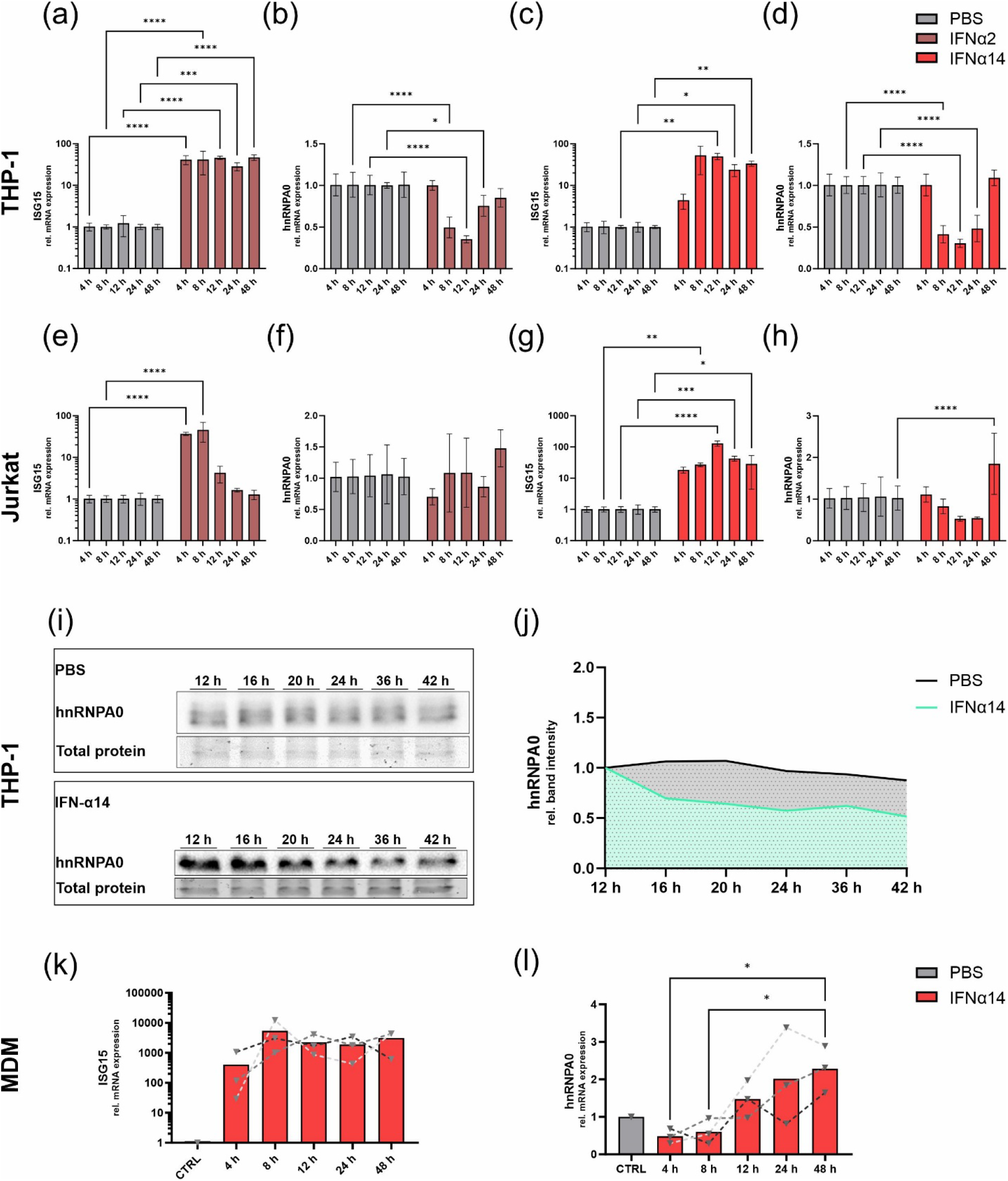
hnRNPA0 mRNA and protein levels are decreased upon IFN stimulation in HIV target cells. Differentiated THP-1 **(a-d)** or Jurkat **(e-h)** cells were treated with IFNα2 (dark red) or IFNα14 (bright red) [10 ng/ml]. At the indicated time points cells were lysed and subjected to the respective read-out. **(a-h)** mRNA expression of **(a,c,e,g)** ISG15 and **(b,d,f,h)** hnRNPA0 in THP-1 and Jurkat cells upon IFN treatment RT-qPCR was performed to monitor expression levels. *ACTB* was used as loading control. **(I,j)** THP-1 cells were treated with 10 ng/ml IFNα14. Proteins were separated by SDS-PAGE, blotted on a nitrocellulose membrane, and analyzed using an antibody against hnRNPA0. Total protein amounts were stained using Trichloroethanol and used for normalization. **(i)** Representative Western blots of hnRNPA0 from THP-1 cells treated with PBS or IFNα14. **(j)** The mean values of the quantification of 4 independent Western blots per condition are shown. **(k,l)** Monocyte derviced macrophages were treated with 10 ng/ml IFNα14. At the indicated time points cells were lysed and RNA was isolated to evaluate mRNA expression levels via RT-qPCR of **(k)** ISG15 and **(l)** hnRNPA0. Expression levels of three independent experiments are shown. **(a-h)** Mean (+SD) is shown of four independent experiments (**(a-d)** at 12-48 h contain three replicates). **(a-j)** Two-way repeated-measures ANOVA with Šidák multiple comparisons test, and **(k,l)** one-way repeated-measures ANOVA with Dunnett post-hoc test were performed to evaluate whether the differences between the groups reached statistical significance (*p<0.05, **p<0.01, ***p<0.001, ****p<0.0001).

Lastly, primary monocyte-derived macrophages (MDMs) from healthy donors were isolated and treated with IFNα14 (Fig. 6k-l). *ISG15* expression was strongly and persistently induced to levels comparable to stimulated THP-1 cells. Notably, following initial repression of hnRNPA0 expression levels in MDMs already 4h after IFNα14 treatment, levels were significantly increased to levels 2-fold higher when compared to the PBS control.

These results further emphasized that hnRNPA0 is differently regulated by IFN-I in a cell-type specific manner.

### hnRNPA0 transcript levels are lower in HIV-1-infected individuals compared to healthy controls

To evaluate whether hnRNPA0 levels might be differently expressed in HIV-1-infected individuals, we isolated PBMCs from HIV-1 acutely and chronically HIV-1-infected individuals and those receiving anti-retroviral therapy (ART) and compared the *ISG15* and hnRNPA0 expression levels to a healthy cohort (Fig. 7). Elevated *ISG15* levels were observed in individuals with an acute and chronic HIV infection, albeit the differences to healthy controls were only significant for the chronically infected cohort (p=0.0246). For ART-treated individuals we observed expression levels for both *ISG15* and *hnRNPA0* that were comparable to the uninfected control cohort. Strikingly, individuals with acute (p=0.0483) and chronic (p=0.0549) HIV-1 infection had lower *hnRNPA0* mRNA levels.

**Figure 7.**
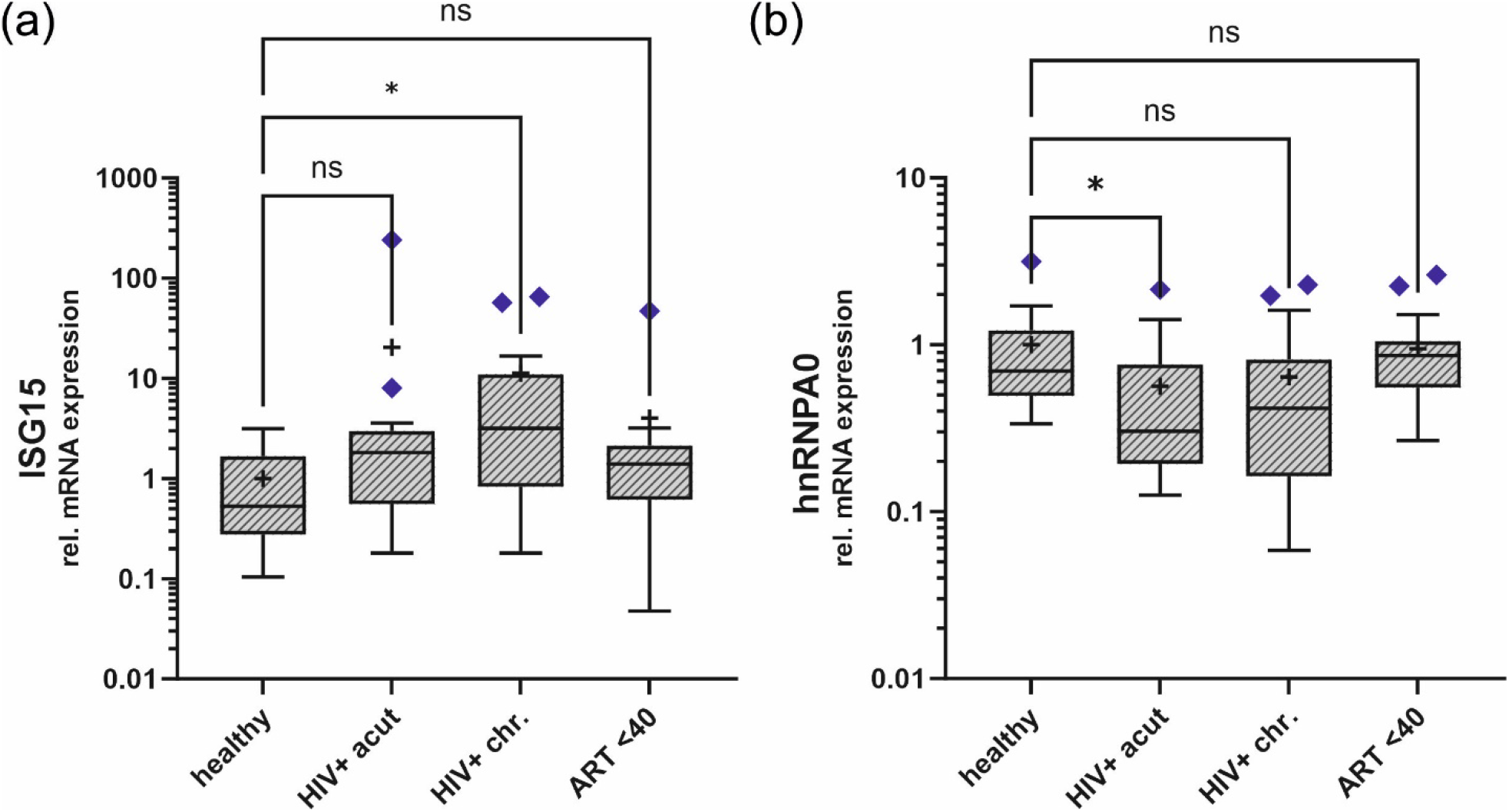
ISG15 and hnRNPA0 mRNA expression levels in HIV-1 infected cohorts. Relative expression levels of **(a)** ISG15 and **(b)** hnRNPA0 of PBMCs of healthy, HIV acutely infected, HIV chronically infected, and HIV infected ART treated individuals. ART <40 patients were below 40 copies/ml (measured by RT-qPCR). ACTB was used for normalization. Mean is indicated as “+”. Error bars are indicated as Tukey min and max values. Purple rectangles represent outliers that were not included into statistical analysis. PMBCs were isolated from 11 healthy donors, 13 with acute HIV-1 infection, 17 treatment-naïve patients with chronic HIV-1 infection and 17 HIV-1 infected patients on ART. Kruskal–Wallis test with the Dunn’s post-hoc multiple comparisons test was used to determine whether the difference between the sample groups reached the level of statistical significance (*p<0.05, and ns, not significant).

In conclusion, the HIV-1-infected therapy-naïve individuals had lower *hnRNPA0* and concomitantly higher *ISG15* levels compared to the healthy controls.

We performed a comprehensive re-analysis of RNA sequencing data obtained from intestinal lamina propria monocyte-derived cells (LPMCs) derived from HIV-1-infected patients (Fig. 8). Our previously published findings demonstrated dysregulation of host-dependent factors, particularly the SRSF family, in these patients (36). Additionally, we observed concurrent upregulation of ISGs, including HIV-1 host restriction factors (54). Compared to other hnRNPs including A1, F or K, which were significantly lower in HIV-1-infected treatment-naïve individuals, hnRNPA0, was rather weakly expressed. However, we detected 1.28-fold lower expression in HIV-1-infected treatment-naïve patients, compared to uninfected controls.

**Figure 8.**
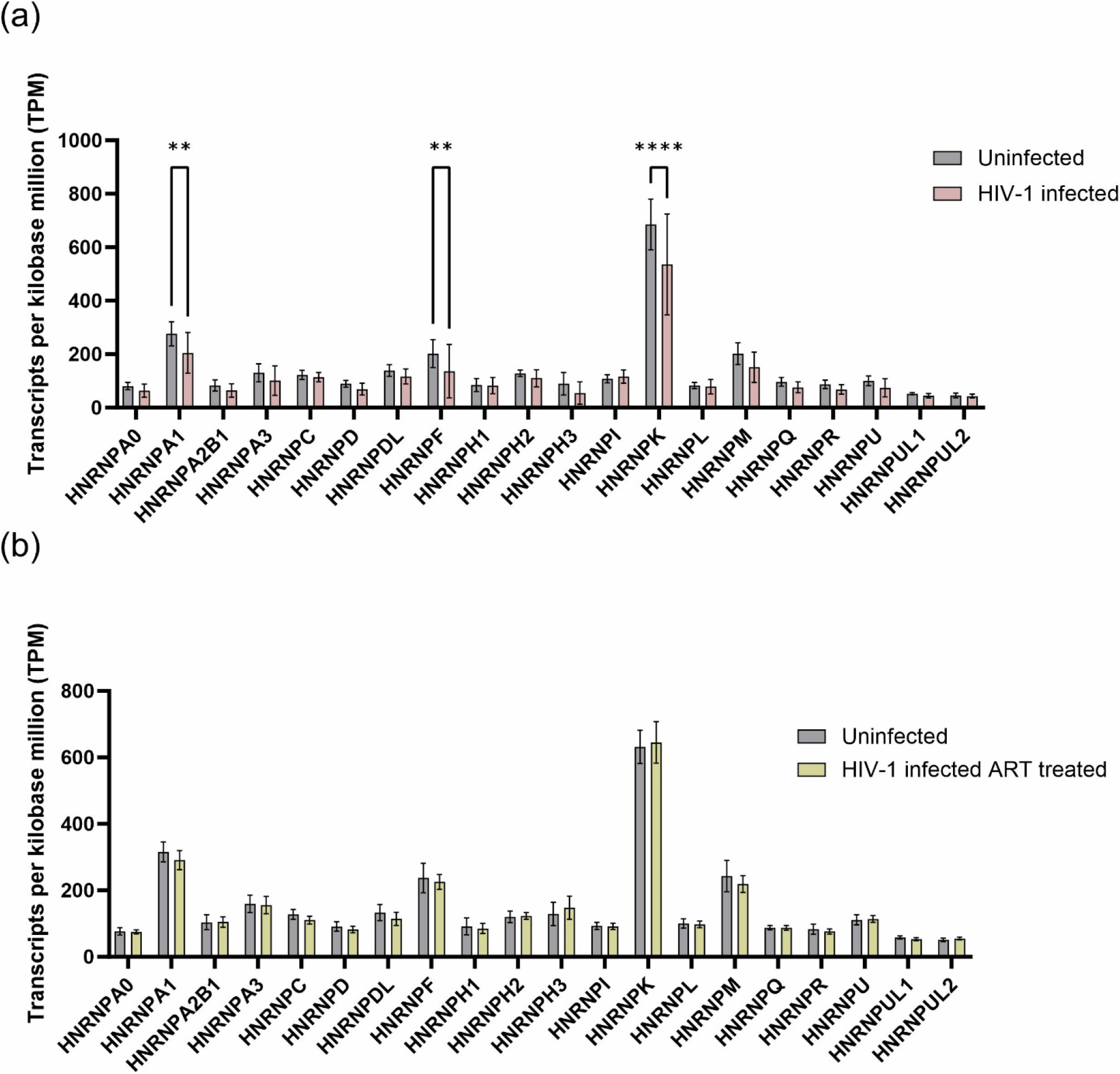
Gene expression levels of hnRNPs in HIV-1-uninfected versus HIV-1-infected and (un-)treated individuals. RNA-sequencing analysis was used to measure transcript levels of hnRNP genes in intestinal lamina propria mononuclear cells (LPMCs). Comparison of the transcript levels of **(a)** HIV-1 infected (n=19) and healthy individuals (n=13) and **(b)** HIV-1 infected ART receiving individuals (n=14) and healthy individuals (n=11). TPM are presented as mean (+ SD). To evaluate whether the differences of the groups reached statistical significance they were compared by two-way ANOVA with Bonferroni post hoc test (*p<0.05, **p<0.01, ***p<0.001 and ****p<0.0001).

In summary, HIV-1 treatment-naïve individuals generally exhibit reduced levels of hnRNPA0 and others in PBMCs and LPMCs.

## Discussion

HIV-1 depends on the host cellular machinery to fully utilize its genome, encompassing processes such as transcription, pre-mRNA splicing, mRNA export, and translation. RNA-binding proteins (RBPs) as members of the heterogeneous nuclear ribonucleoproteins (hnRNP) and serine-arginine-rich splicing factor (SRSF) families play indispensable roles in HIV-1 replication. Various RBPs from these families have been characterized for their interactions with HIV-1 and their functions as host-dependent factors (HDFs).

Mechanistically, RBPs were described to positively or negatively influence the transcriptional efficiency of HIV-1 by interacting with the viral LTR promoter region affecting the recruitment of transcriptional regulators (55–57) and thereby influencing the overall production of viral RNAs (36). Furthermore, during HIV-1 replication multiple spliced and unspliced mRNA transcripts which are categorized into 2, 4 and 9kb classes, are synthesized. hnRNPs are involved in regulating the HIV-1 alternative splice site usage, which determines the production of different mRNA isoforms coding for viral proteins. When hnRNPA0 levels are elevated *in vitro* we observed a drastic decrease in viral particle production and viral copies in the supernatant, which led to a significant decrease in infected TZM-bl reporter cells (Fig. 1f-i), exposing a potent antiviral activity.

This antiviral activity was not due to inefficient alternative splice site usage, the predominant effect reported in similar studies with multiple other hnRNPs (47, 58–61), but rather due to a decrease in LTR-activity, which resulted primarily in decreased total viral mRNA (exon 1 and exon 7; Fig. 2b). *In silico* mapping of hnRNPA0 motifs revealed multiple binding sites in the HIV-1 genome (Supp.Fig.1). Despite not obtaining the highest Z-scores (max. 3.18) several binding sites are located within the HIV-1 LTR (Supp.Fig.2).

hnRNPs might interact with other nucleic-acid-binding proteins either via their auxiliary domain, which in case of hnRNPA1 is the unstructured glycine-rich domain (62, 63), or via the RBD in case of hnRNPA2B1 (12). Therefore, hnRNPA0 could act as mediator facilitating LTR transcription by binding or scaffolding of other nucleic-acid related proteins, already observed for other hnRNPs (64–68). Additionally, hnRNPA0 could potentially interact with the 7SK particle, which binds and thus inhibits the function of P-TEFb (69) crucial for HIV-1 transcriptional elongation (70). Notably, knockdown of hnRNPA1 and A2B1 (71, 72) attenuated the dissociation of P-TEFb from the 7SK complex, resulting more active P-TEFb. However, based on our results, hnRNPA0 would rather facilitate the inhibition i.e. binding of P-TEFb by the 7SK complex. Although we cannot exclude the mentioned possibilities, it seems more reasonable that hnRNPA0 might directly bind to the LTR as we observed a decrease in LTR-activity under depleted hnRNPA0 levels and in the absence of Tat (Supp.Fig.4). By using electrophoresis mobility shift assays (EMSA) DNA-binding capacity has been observed for several hnRNPs, including hnRNPA1 (73) and A3 (74), the latter one binding DNA via its RNA-recognition-motif 1 (RRM1) domain. Further, hnRNPK (75–80), and U were not only able to bind DNA, but were also directly involved in transcription (81), with hnRNP K primary known as a transcription factor. Furthermore, it has been reported that hnRNPA1 and A2B1 can exert opposing effects on HIV-1 particle production and infectivity, with repression under high expression levels and enhancement under low expression levels. The observed effects were predominantly attributed to their impact on viral transcriptional activity. Importantly, hnRNPA3 showed no significant effect on viral replication, highlighting the selective role of hnRNPA/B family members in HIV-1 gene regulation (55). Furthermore, a competitive binding interaction, which resulted in the repression of the LTR activity when Tat was present, was already observed for the RBP SRSF1 by binding to overlapping sequences within TAR and the 7SK RNA (56). Notably, SRSF1 also increased the basal level of viral transcription when Tat was absent, a comparable observation was made for hnRNPA0 in our study (Supp.Fig.4). The hypothesis that hnRNPA0 competitively binds to the LTR is further supported by the increased LTR activity of the siRNA transfected reporter cells (Fig. 2l). However, further studies are needed to elucidate the mode of action, but based on our findings, hnRNPA0 emerges as a promising candidate for shock-and-kill or block-and-lock follow up studies, given its ability to modulate LTR activity depending on its expression levels.

During PRF in the course of translation the ribosome shifts the viral reading frames (−1) at specific RNA secondary site in the *gag-pol* overlap region, enabling the production of the Gag-Pol polyprotein (82). HIV-1 (83) uses -1PRF to maintain correct stoichiometry of Gag and Pol proteins, which are essential for virion assembly and maturation. During this study, we demonstrated that hnRNPA0 might significantly regulate viral frameshifting in HIV-1, although we cannot discriminate between a direct or indirect interaction. Previously described proteins regulating viral -1PRF are among others zinc finger antiviral protein (ZAP) (84) and the interferon-stimulated gene C19Orf66 (Shiftless, SFL) (85). Of note, in SARS-CoV-2 context, hnRNPH1 and H2 were able to reduce -1PRF in an intensity comparable to ZAP (84). Interacting with the ribosome requires a cytoplasmic localization of the RBPs. It has already been observed, that hnRNPA1 and A2B1 are capable of shuttling between the cytoplasm and the nucleus (12, 86). Given that hnRNPA0 has been identified both as a nuclear and cytoplasmic RBP in neurons, it is plausible that it possesses the capability for nucleocytoplasmic shuttling (87).

Studies have demonstrated that the binding affinity between SFL and its target RNA can be influenced by RBPs (88). Consequently, it is intriguing to hypothesize that hnRNPA0 may act as a chaperone for SFL, facilitating its association with HIV-1 mRNA and thereby promoting frameshift inhibition.

Interferons (IFNs) play a crucial and indispensable role in the host’s defense against viral infections. They are essential components of the innate immune response, being rapidly induced upon viral recognition (45, 46). IFNs activate a complex signaling cascade leading to the reprogramming of the cellular expression profile, by inducing interferon-stimulated-genes (ISGs) and repressing interferon-repressed-genes (IRepGs), which collectively establish an antiviral state inhibiting viral replication, limiting viral spread, and enhancing overall antiviral immunity (89, 90).

Previously we were able to show that the expression of SRSF1, an essential RBP and host dependency factor is modulated by IFN-I (36). In this study, we identified hnRNPA0 as interferon-regulated gene, which is directly or indirectly regulated by a JAK1/2 dependent pathway. In fact, JAK1 can phosphorylate STAT1 (91) and STAT1 is defined as a transcription factor of hnRNPA0 by the *ENCODE Transcription Factor Targets* data set (92, 93). Which mechanism might facilitate a repression in mRNA levels of hnRNPA0 remains unclear, however it is plausible that STAT1 promotes the substantial transcription of hnRNPA0 mRNA, which subsequently underlies auto-regulation similar to what has been previously reported for various other hnRNPs (94–98). With hnRNPA0 preferably binding to AU-rich elements (AREs), which are predominantly located in the 3’ UTR, it is most likely that hnRNPA0 binds a regulatory element within its 3’ UTR and either inhibits or causes premature polyadenylation of its own mRNA (99). Another regulatory level involves IFN-induced phosphorylation, since hnRNPA0 can be phosphorylated by MAPK-activated protein kinases (MAPKAPKs) 2 and 3 (100). Particularly, MAPKAPK2 has been demonstrated to phosphorylate hnRNPA0 at Serine 84 residue in response to Lipopolysaccharide (LPS) stimulation. Additionally, hnRNPA0 is regulated by short RNAs complementary to ribosomal 5.8S RNA (101) and miR205HG that has been identified as an translational inhibitor (102). Nevertheless, further investigations are needed to elucidate the mechanisms underlying hnRNPA0 regulation upon IFN-treatment.

Treating HIV-1 target cells with IFNα2 and IFNα14 revealed cell-type and IFN-specific differences with macrophages showing the strongest effects. Interestingly, THP-1 cells and MDMs differed in their expression patterns as an overexpression of hnRNPA0 was observed in MDMs but not in THP-1 cells, even when we additionally monitored later time points (data not shown).

Concomitantly to high ISG15 mRNA expression, we observed low hnRNPA0 mRNA levels in PBMCs isolated from HIV-1 acutely and chronically infected individuals, however, the impact of hnRNPA0 downregulation induced by interferons, even if temporarily limited, remains unclear in the *in vivo* context. Depleted hnRNPA0 levels significantly increased the LTR-activity and consequently resulted in higher production of infectious viral particles (Fig. 4). Whether this plays a facilitating role for virus replication during acute infection cannot be answered at this point and requires further studies. Consistent with prior studies (103), the expression levels of hnRNPA0 in ART-treated patients were found to be comparable to those in the naïve cohort. This observation suggests that only ongoing viral replication may result in decreased hnRNPA0 levels due to heightened Jak1/2 induction (104), thereby partially counteracting or compensating for the direct antiviral effects exerted by ISGs and restriction factors by higher LTR activity.

This study has limitations: hnRNPs might interact with the viral RNA genome and facilitate its packaging into new viral particles during the assembly process. hnRNPs can impact the stability of HIV-1 RNA by binding to specific regions within the viral transcript and influencing its degradation or stabilization. No packaging and stability studies were performed in this study and future work is needed to address these issues. A possible influence of hnRNPA0 on the expression of host immune factors, including interferons and cytokines, affecting the host’s ability to control HIV-1 infection, was not analyzed. hnRNPs participate in the transport of HIV-1 RNA from the nucleus to the cytoplasm, where viral translation and assembly occur. Since in this study the RNA localization was not analyzed, we cannot exclude a direct or indirect effect of RNA trafficking as another level of inhibition by hnRNPA0. No multiround replication experiments were performed since long term overexpression of hnRNPA0 was technically challenging due to autoregulatory mechanisms as discussed above. The regulation of hnRNPA0 mRNA levels remains to be further analyzed in the context of multiple cell types, in particular primary cells. Given the initial repression, which turned into an overexpression in Jurkat T-cells and MDMs a timeline of hnRNPA0 levels connected to HIV-1 pre and post infection states in different cell-types would also be reasonable. Although it is very likely that hnRNPA0 binds the HIV-1 LTR, further studies are needed to prove a direct binding of hnRNPA0 to the 5’ LTR sequence.

Additional studies are also required to elucidate the mechanistic role of hnRNPA0 in frameshifting.

## Material and methods

### Cell culture and preparation of virus stocks

PBMC isolation procedures, cell culturing and preparation of virus stocks were performed as described elsewhere (105).

### Transient transfection and siRNA-mediated knockdown

Vero cells seeded into 12-well plates and incubated overnight were transfected using plasmids harboring different HIV-1 LTRs upstream of a Firefly luciferase, and plasmids encoding HIV-1 Tat (SVcTat) (106) and FLAG-tagged hnRNPA0 (pcDNA3.1-FLAG-NLS-hnRNPA0). A promoter-lacking but luciferase encoding plasmid was used as a control (pTA-Luc).

A549-LTR-Luc-PEST reporter cells were generated using the Sleeping Beauty transposase system (43, 107). For the LTR-assay, 24h post seeding cells were transfected with siRNA targeting hnRNPA0 (s21545 (Thermo Scientific)) and a plasmid encoding HIV-1 Tat (SVcTat). Off-target siRNA (Silencer Select Negative Control #2, Thermo Scientific), and an empty vector control (pcDNA3.1).

24h (Vero) or 48h (A549-LTR-Luc-PEST) post transfection cells were lysed using Lysis-Juice (PJK), incubated for 15min under agitation subjected to a freeze and thaw cycle. Lysates were centrifuged for 10min at 13,000rpm using a tabletop centrifuge and transferred into an Immuno 96 MicroWell plate (Nunc). Luciferase assays were performed using the GloMax Discover (Promega) and Beetle-Juice (PJK) or Luciferase Assay System (Promega).

To perform dual luciferase frameshift assays HEK293T cells were seeded 24h prior transfection in 12-well plates. Cells were transfected with varying amounts of pcDNA3.1 FLAG NLS hnRNPA0. After 6h the cells were transfected with the reporter plasmid pDual-HIV either containing the wildtype frameshift site with Renilla luciferase in frame and Firefly luciferase in the -1 frame or the mutated frameshift site with both luciferases in frame. After 24h the cells were harvested using protein lysis buffer (Promega) and centrifuged for 10min at 13,000rpm. Lysates were transferred into a white 96-well assay plate. Luciferase reporter assays were performed on the GloMax Discover (Promega) using the Dual Luciferase Reporter Assay System (Promega). Frameshift efficiency was determined by the ratio of Firefly luciferase to Renilla luciferase.

### Detection of cellular and viral RNA, proteins and HIV-1 infectivity

RNA isolation, quantitative and semi-quantitative RT-PCR and p24-CA ELISA were performed as described previously (105). Primary antibodies used in this study are listed in Table 1. The workflow to determine viral infectivity using TZM-bl cells is described elsewhere (47, 105). Detection

**Table 1.** Primary antibodies used in this study.

| Target | Supplier |
| --- | --- |
| hnRNPA0 | Abcam (ab197023) |
| FLAG-tag | Abcam (ab49763) |
| HIV-1 Vif | Abcam (ab66643) |
| HIV-1 p24 | Aalto (D7320) |
| APOBEC3G | HIV Reagent Program |
| HIV-1 p15 | Abcam (66951) |
| HIV-1 Vpr | Proteintech (51143-1-AP) |
| Human CD11b | Biolegend (301306) |

### Statistical analysis

Differences between two groups were analyzed by unpaired two-tailed student’s or Welch’s t-test. Multiple group analyses were performed using one- or two-way ANOVA followed by Bonferroni, Dunnett’s or Tukey’s *post hoc* test. Mixed-models followed by Dunnett’s *post hoc* tests were used for time series analysis of multiple groups. A Kruskal-Wallis test with the Dunn’s post hoc multiple comparisons test was applied to compare mRNA levels in PBMCs from acutely and chronically HIV-1-infected patients as well as from healthy donors due to violation of the assumptions for a parametric test. Outlier were identified via Tukey’s range test and excluded from further statistical analysis. If not indicated differently, all experiments were repeated in three independent replicates. Asterisks indicated p-values as *p<0.05), **p<0.01), ***p<0.005) and ****p<0.0001).

## Acknowledgements

We thank Christiane Pallas and Barbara Reimer for excellent technical assistance. We thank Heiner Schaal for providing plasmid DNA. The following reagents were obtained through the AIDS Research and Reference Reagent Program, Division of AIDS, NIAID, NIH: Panel of Full-Length Transmitted/Founder (T/F) Human Immunodeficiency Virus Type 1 (HIV-1) Infectious Molecular Clones, HRP-11919 contributed by Dr. John C. Kappes. TZM-bl cells from Dr. John C. Kappes and Dr. Xiaoyun Wu. ApoC17 antibody from Dr. Klaus Strebel.

## Ethic statement

This study has been approved by the Ethics Committee of the Medical Faculty of the University of Duisburg-Essen (14-6155-BO, 16-7016-BO, 19-8909-BO). Form of consent was not obtained since the data was analyzed anonymously. The funders had no role in study design, data collection and analysis, decision to publish, or preparation of the manuscript.

## Data availability

Next-generation sequencing data were deposited at the NCBI Sequence Archive Bioproject PRJNA422935. Further inquiries can be directed to the corresponding author

## Supplementary Material

**Supplementary Figure 1.**
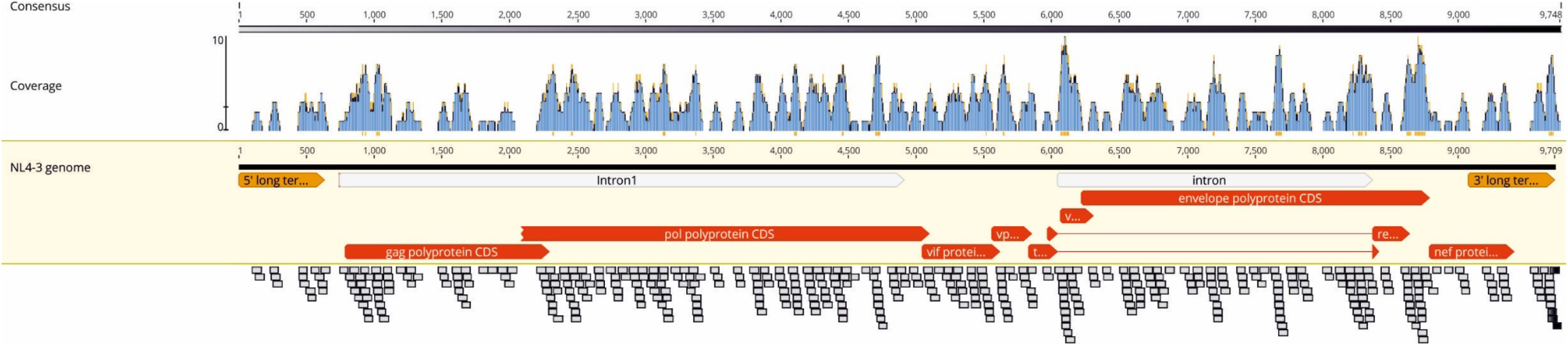
hnRNPA0 binding sites within the subviral clone NL4-3. The binding sequence motifs of hnRNPA0 (108) were mapped to the subviral clone NL4-3 using RBPmap (109). Regions with more than 6 clustered binding sites are highlighted in orange below the coverage graph. The frequency of binding sites per region is indicated by the coverage, shown in blue bars. The respective sequences are shown below the NL4-3 genome.

**Supplementary Figure 2.**
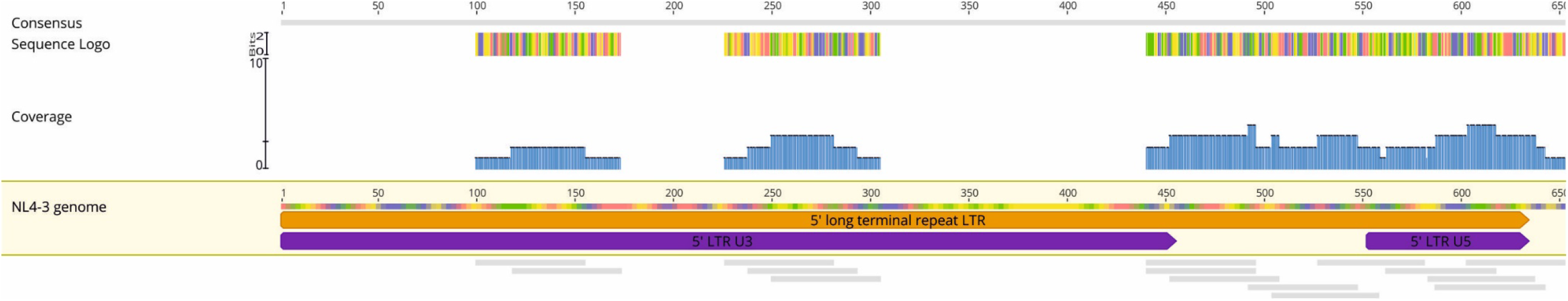
hnRNPA0 binding sites within the NL4-3 5’ LTR. The binding sequence motifs of hnRNPA0 (108) were mapped to the subviral clone NL4-3 using RBPmap (109). The frequency of binding sites per region is indicated by the coverage, shown in blue bars. The respective sequences are shown below the NL4-3 genome. The unique 3 (U3) and unique 5 (U5) elements of the 5’ LTR promoter are indicated in purple.

**Supplementary Figure 3.**
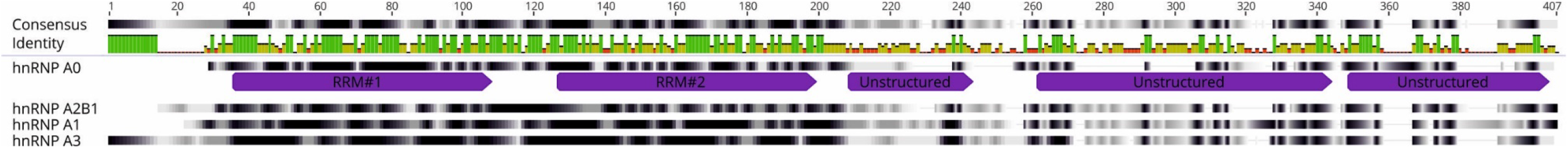
Protein alignment of the four members of the hnRNP A/B family. Clustal Omega alignment of the canonical sequences of hnRNPA0, A1, A2B1, A3. The similarity of sequences was calculated using a Blosum62 score matrix and visualized using different shades of grey with black indicating 100% similarity. Mean pairwise identity over all pairs in the column is visualized under “Identity” using bars with a color code from red to green with tall green bars indicating 100% similarity between the pairs in the respective region. The protein sequences were ordered according to similarity to each other. Sequences were obtained from the Uniprot database (33) with the respective entry IDs: hnRNPA0: Q13151, hnRNPA1: P09651, hnRNPA2B1: P22626, hnRNPA3: P51991. Domains of hnRNPA0 were annotated according to SMART prediction (110). RRM = RNA-recognition-motif

**Supplementary Figure 4.**
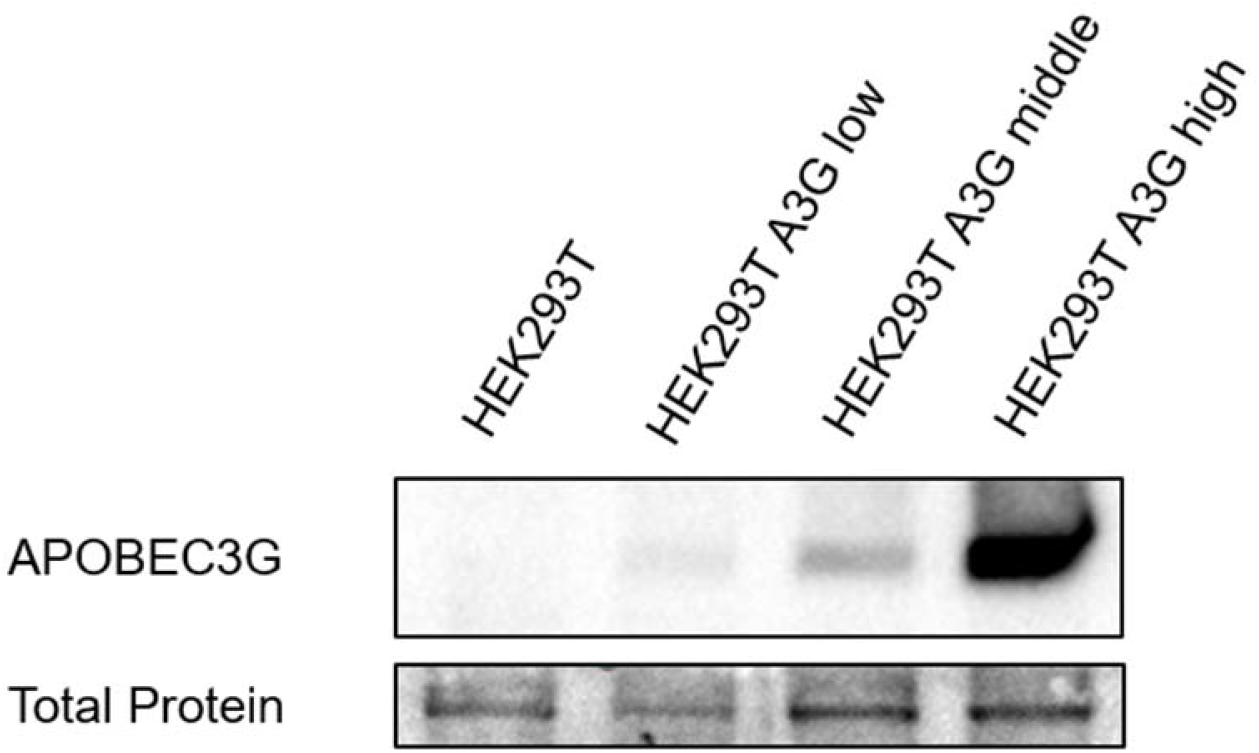
A3G protein levels of HEK293T A3G expressing cells. HEK293T were previously stably transfected with a plasmid constitutively expressing APOBEC3G (A3G) and sorted in A3G low, middle and high expressing cells. Cells were cultured for 24h before being washed with PBS and lysed. Proteins were isolated, separated via PAGE and A3G amounts were analyzed via Western blotting.

**Supplementary Figure 5.**
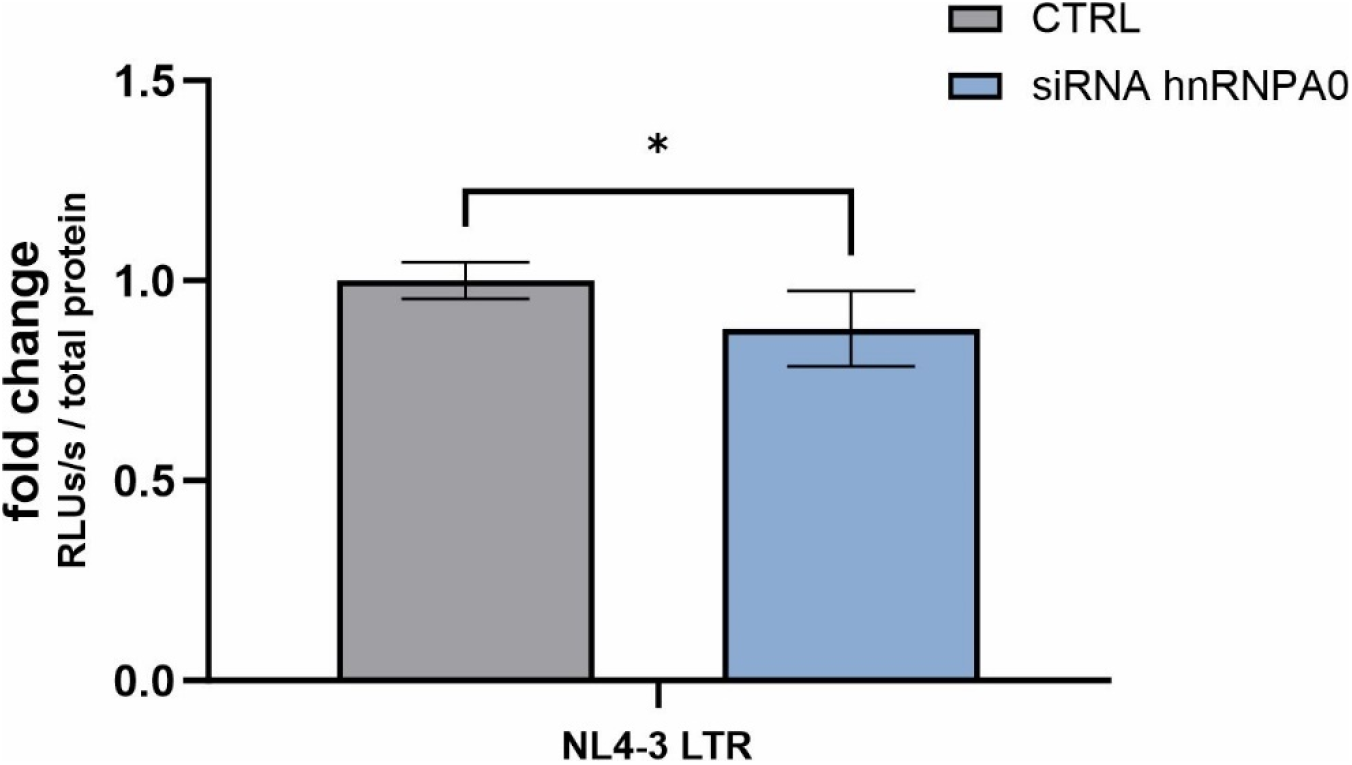
NL4-3 LTR activity is decreased under depleted hnRNPA0 levels in the absence of Tat. A549 LTR Luc-PEST cells were transfected with either siRNA targeting hnRNPA0 or a non-template siRNA as control. 48h post transfection cells were rinsed with PBS, lysed and the luciferase activity was measured via luciferase assay. Unpaired two-tailed t-tests were calculated to determine whether the difference between sample groups reached the level of statistical significance (*p<0.05, **p<0.01, ***p<0.001, ****p< 0.0001 and ns, not significant)

**Supplementary Figure 6.**
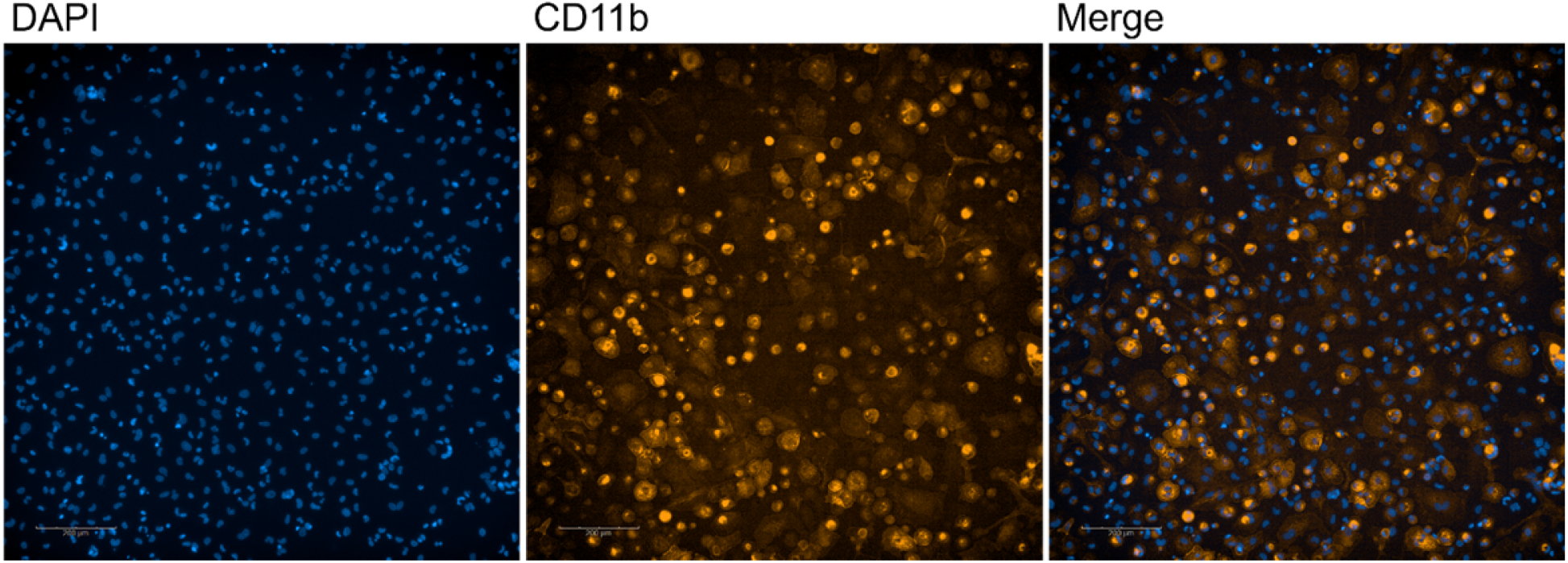
CD11b staining in differentiated THP-1 cells. THP-1 cells were seeded into 6-well plates and treated with 100nM Phorbol-12-myristat-13-acetat (PMA). 5 days post treatment cells were washed using PBS, fixed for 10min using 3% formaldehyde and permeabilized for 10min using 0.1% Triton X-100. Unspecific binding sites were blocked using 2% BSA for 20min before immunohistology staining was performed. Nuclei were stained using DAPI and CD11b was stained as a marker for differentiation using an PE labeled antibody directed against CD11b.

